# Honeybees optimize their foraging behaviour in relation to spatio-temporal changes in nectar and pollen availability

**DOI:** 10.1101/2020.07.08.193268

**Authors:** Jan J. Kreider, Anna Nehrkorn, Svenja Bänsch, Carmen Kirsch, Catrin Westphal

**Author notes:** Corresponding author: J.J. Kreider. Equally contributing first authors (Jan J. Kreider, Anna Nehrkorn).

## Abstract

Intensified agriculture increasingly threatens wild and managed bees by promoting landscape uniformity and reducing floral resource availability whereas urban areas can provide continuous floral resources within green spaces and private gardens. Mass-flowering events of crops and trees, such as lime trees (*Tilia* spp.), can provide ample floral resources but only for short time periods. Using waggle dance decoding, pollen analysis and bee abundance recordings, we investigated the temporal shift in honeybee foraging behaviour in response to lime tree mass-flowering. Honeybees in urban areas extended their foraging range during lime tree flowering. Foraging behaviour of honeybees in rural areas did not change to such an extent and honeybees foraged in sown flower strips. Our results suggest that honeybees optimize their foraging behaviour to exploit highly rewarding resources instead of extending foraging ranges in times of floral resource scarcity.

## 1. Introduction

Bees are increasingly threatened by a variety of stressors such as parasites, pesticides, habitat loss and food scarcity resulting in declined bee abundance and bee species diversity (Brown and Paxton 2009; Goulson et al. 2015; Cariveau and Winfree 2015). As bees are important pollinators of agricultural crops, threats to bees can put agricultural yields and global food security at risk (Potts et al. 2010; Garibaldi et al. 2013). Bee declines are particularly apparent in areas with intensive agriculture which are typically uniformly structured, do not constantly provide flowering plants throughout the entire season and contain large proportions of non-entomophilous crops (Steffan-Dewenter and Kuhn 2003; Kovács-Hostyánszki et al. 2017). Urban areas, in contrast, provide temporarily constant flower resources and higher flower diversity (e.g. in green spaces and private gardens), resulting in higher bee abundance and bee species diversity (Theodorou et al. 2016; Martins et al. 2017). Temporary floral resource scarcity in rural areas might especially be costly for agricultural production as continuous resource availability is crucial to support pollinators, and hence the delivery of ecosystem services (Schellhorn et al. 2015). Low floral resource availability has been shown to negatively affect size and health of colonies of the European honeybee (*Apis mellifera* L.), the most important pollinator worldwide, making them more prone to damages by the parasitic mite *Varroa destructor* Anderson & Trueman (Brodschneider and Crailsheim 2010; Requier et al. 2017; Wintermantel et al. 2019).

Resources are not only variable in space and time, but also differently used by bees depending on functional traits. Accordingly, bees can be grouped into honeybees, bumblebees and other wild bees (from here on referred to as wild bees; Rollin et al. 2013). Wild bees comprise mainly solitary species, use locally available flower resources in semi-natural habitats (SNH) and some species are specialized to feed on particular plant families, genera or species (Michener 2000; Rollin et al. 2013). Bumblebees form independently-founded annual colonies composed of 70 to 1800 individuals (Cueva del Castillo et al. 2015) and use both locally available resources in SNH and highly rewarding resources, such as agricultural crops or flower strips (Westphal et al. 2006a; Rollin et al. 2013). Most bumblebee species forage on a vast variety of flower species. Their foraging distances can reach up to 3000 m (Westphal et al. 2006b; Wolf and Moritz 2008) and foraging behaviour is adapted to spatio-temporal availability of floral resources (Westphal et al. 2006a). Honeybees live in colonies of several thousand individuals with queens that reproduce over multiple seasons and hibernate as a colony. They mostly forage for highly rewarding resources and due to their complex social organization foraging sites can be communicated to nestmates (Rollin et al. 2013; Sponsler et al. 2017). Foraging distances of honeybees are often below 1000 m, but in extreme cases foraging distances can reach up to 14 km (Visscher and Seeley 1982; Beekman and Ratnieks 2000; Couvillon et al. 2014; Danner et al. 2016; Balfour and Ratnieks 2017; Danner et al. 2017; Bänsch et al. 2020).

If floral resources are locally scarce honeybees uphold the required food quality and quantity by increasing their foraging range (Steffan-Dewenter and Kuhn 2003; Danner et al. 2016; Danner et al. 2017; Bänsch et al. 2020). After mass-flowering events of agricultural crops (such as oil seed rape or sunflowers) honeybees in rural landscapes switch to flower resources in SNH or to agricultural weeds (Rollin et al. 2013; Requier et al. 2015; Sponsler et al. 2017). Pollen resource use by honeybees can be inferred from pollen load analysis and nectar resource use from honey stomach analysis of returning foragers (Garbuzov et al. 2015; Danner et al. 2016; Balfour and Ratnieks 2017; Marzinzig et al. 2018; Bänsch et al. 2020).

Honeybees communicate the distance and direction to a rewarding foraging site to their nestmates with a waggle dance which is performed by returning foragers (von Frisch 1967; Couvillon 2012). Waggle dance decoding can be used to detect foraging habitats of honeybees, however identifying exact foraging locations is impossible (Schürch et al. 2013; Balfour et al. 2015; Balfour and Ratnieks 2017; Bänsch et al. 2020).

Here we investigate the spatio-temporal changes of honeybee foraging behaviour in rural and urban landscapes in response to lime tree (*Tilia* spp. L.) mass-flowering. We inferred honeybee foraging habitats from waggle dance decoding and analysed resource use by pollen analysis. Additionally, we determined abundances of honeybees, bumblebees and wild bees in foraging habitats. We addressed the following hypotheses: (1) According to their foraging strategies honeybees, bumblebees and wild bees exploit different foraging habitats. (2) Mass-flowering trees and sown flower strips are the most important foraging resources for honeybees in urban and rural areas, respectively. (3) Honeybees tend to exploit highly rewarding resources and adapt their foraging distances to spatial and temporal variation in relation to the availability of floral resources in different foraging habitats.

## 2. Material and Methods

### Experimental design

We positioned two honeybee observation hives each in an urban landscape (41% urban area within 1 km radius) of Göttingen (Lower Saxony, Germany) and in a rural landscape (94% arable field within 1 km radius) south of the city. Each observation hive had two frames positioned above each other and consisted of approximately 2000 worker bees and one queen bee (Supplement Fig. 1). In the field we mapped the most dominant land use types and potential bee foraging habitats within a radius of 1 km around the locations of the hives using QGIS 3.12.3 and classified them as arable field, sown flower strip (mostly *Phacelia tanacetifolia* Benth.), SNH (including forests, extensively used grasslands and botanical gardens) and urban area (including built-up areas, roads, green spaces and private gardens). Additionally, we mapped aggregations of more than two lime trees.

**Fig. 1.**
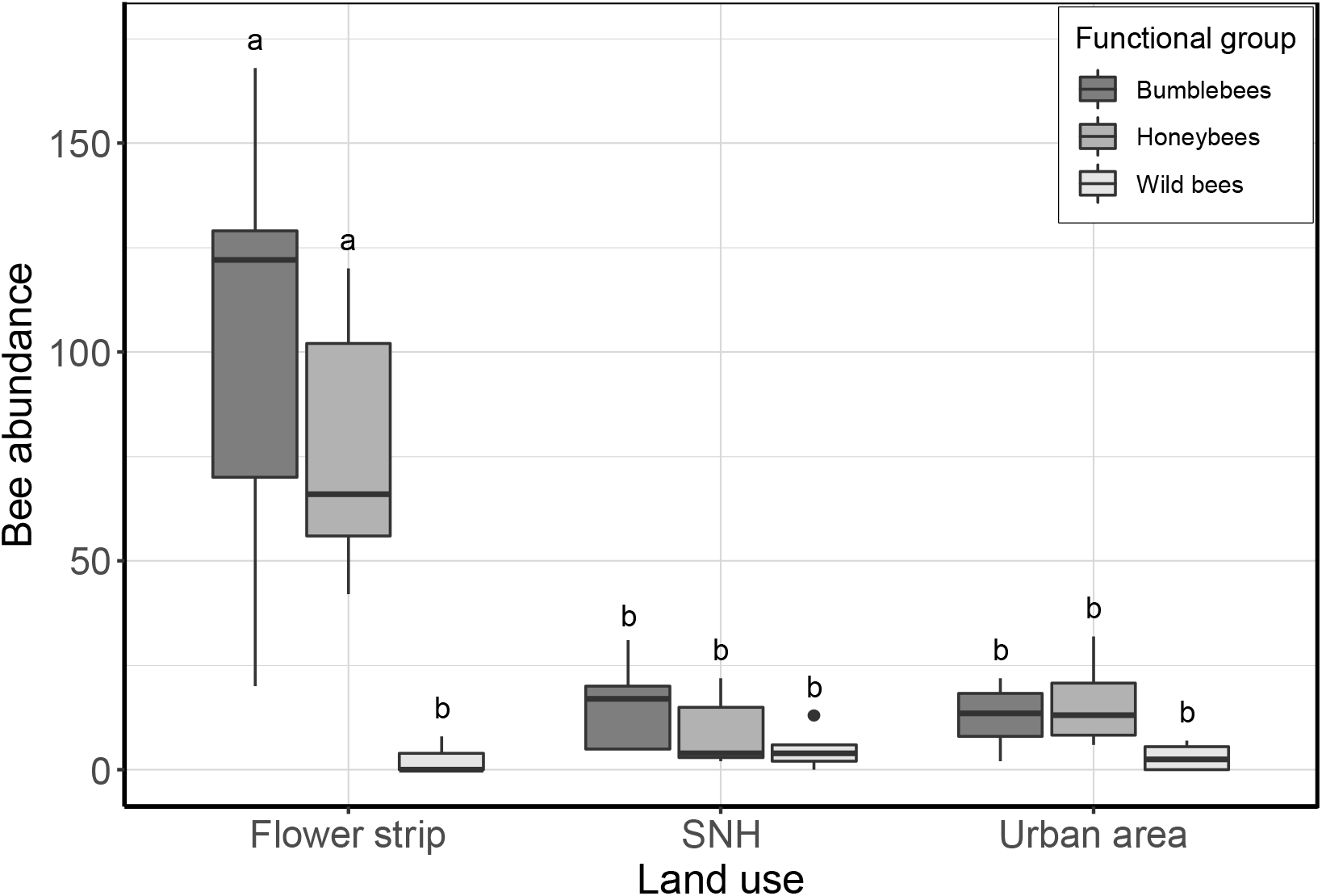
Abundance of bumblebees, honeybees and wild bees in flower strips, semi-natural habitats (SNH) and urban areas. Letters over boxplots show differences between groups from Tukey’s pairwise post-hoc test (α = 0.05). GLMM results in Table I. Pairwise comparisons in Supplement Table I.

### Waggle dance decoding

During June and July 2018, we recorded honeybee waggle dances of returning foragers over a period of four weeks resulting in four record days per hive of which two were during and two after lime tree flowering. Records were taken at random times of the day under similar weather conditions with temperatures above 15°C, no rain and little wind. Video recordings of the bee activity in the hives lasted for 30 minutes and were conducted using a camcorder (Sony HDR CX240E). We decoded all waggle dances of nectar and pollen foragers with four or more circuits following the protocol from Couvillon et al. (2012). If dances consisted of more than four circuits, we decoded four consecutive circuits in the middle of the dance leaving out the first and last circuit. The angle of the dance was measured in relation to a vertical line to the ground. The direction of the dance can be decoded by adding the angle of the dance to the vertical line to the azimuth of the sun at the time of the dance. From the four decoded circuits, we calculated the mean duration and the mean direction of the waggles. All dances which aimed at foraging sites closer than 100 m were considered to be local dances and the distance to the foraging site was set to 100 m as waggle dance decoding can be erroneous for such dances (Schürch et al. 2013).

### Foraging habitats in urban and rural areas

On each observation date, we decoded the first observed dance per hive in the field. We estimated the distance and direction to the foraging location and chose a flower-rich habitat which presumably was the foraging site of the dancing honeybee. Within those habitats we conducted 5 min standardized transect walks during which we counted honeybees, bumblebees and wild bees and classified the habitats as sown flower strip, SNH (transects in hedgerows and field margins) or urban area (transects in green spaces and gardens). After decoding all waggle dances, we descriptively analysed the spatial distribution of foraging habitats within the area around the hives in which we had mapped land use.

### Pollen analysis

During lime tree flowering we sampled three honeybees with pollen loads and three honeybees without pollen loads from each hive assuming them to be pollen and nectar foragers, respectively. Overall, we obtained pollen loads from the legs of twelve pollen-collecting honeybees and pollen from the honey stomachs of twelve nectar-collecting honeybees. We identified the amount of lime tree and *P. tanacetifolia* pollen within approximately 200 pollen grains per sampled honeybee. If less than 200 pollen grains were present in the sample, we identified lime tree and *P. tanacetifolia* amount among all pollen grains. *P. tanacetifolia* was the main flowering plant in flower strips and does not grow naturally in the study area. As the number of pollen grains counted differed between samples of foraging honeybees, we first calculated the proportion of lime tree and *P. tanacetifolia* pollen grains within samples from individual honeybees. Subsequently, we calculated the mean proportion of lime tree and *P. tanacetifolia* pollen from pollen loads and from honey stomachs each from the urban and the rural landscape.

### Data analysis

We compared abundances of the three functional bee groups (bumblebee, honeybee, wild bee) in different transect habitats (flower strip, SNH, urban area) using generalized linear mixed models (GLMM) with a gaussian distribution and square-root-transformed response variable to satisfy model assumptions using the function *glmmTMB* from the R package ‘glmmTMB’ (Brooks et al. 2017). Landscape (urban, rural) and a transect identifier were included as nested random term.

We analysed foraging distances in the different landscapes (urban, rural) during and after lime tree flowering with linear mixed-effects models (LME) with the function *lme* from the R package ‘nlme’ (Pinheiro et al. 2018). As we conducted repeated observations on the same hives, a hive and record identifier were included as nested random term.

For both models, we performed Tukey’s pairwise post-hoc test with a significance level of 0.05 using the function *emmeans* from the R package ‘emmeans’ (Lenth 2018). All statistics were performed in R 3.5.0 (R Core Team 2018).

## 3. Results

### Bee abundance in foraging habitats

In total, we conducted transects in seven flower strips (FS), in five SNH and in four urban areas (UA) and recorded 837 bumblebees (B), 654 honeybees (H) and 53 wild bees (W). Bumblebee and honeybee abundances in flower strips were significantly higher than wild bee abundance in any habitat type and higher than bumblebee and honeybee abundances in SNH and urban areas (mean abundance ± SE; FS: B 101.1 ± 19.0, H 77.7 ± 11.3, W 2.3 ± 1.2; SNH: B 15.6 ± 4.9, H 9.2 ± 4.0, W 5.0 ± 2.2; UA: B 12.8 ± 4.3, H 16.0 ± 5.8, W 3.0 ± 1.8; GLMM results in Table I; pairwise comparisons in Supplement Table I; Fig. 1).

**Table I.** Generalized linear mixed model (GLMM) results of square-root-transformed abundance of functional bee groups (bumblebee, honeybee, wild bee) in different transect habitats (flower strip, SNH, urban area). GLMM was fitted with a gaussian distribution. Pairwise comparisons in Supplement Table I.

|  | **Bee abundance** |  |  |  |
| --- | --- | --- | --- | --- |
|  | Estimate | SE | z | p |
| Intercept | 9.70 | 0.59 | 16.32 | <0.001 |
| Honeybee | -1.02 | 0.84 | -1.22 | 0.223 |
| Wild bee | -8.72 | 0.84 | -10.38 | <0.001 |
| SNH | -5.97 | 0.92 | -6.48 | <0.001 |
| Urban area | -6.35 | 0.99 | -6.44 | <0.001 |
| Honeybee:SNH | 0.04 | 1.30 | 0.03 | 0.976 |
| Wild bee:SNH | 6.89 | 1.30 | 5.29 | <0.001 |
| Honeybee:Urban area | 1.48 | 1.39 | 1.07 | 0.287 |
| Wild bee:Urban area | 6.60 | 1.39 | 4.73 | <0.001 |

### Waggle dance decoding

In total, we decoded 165 waggle dances with a mean foraging distance of 387.5 m (±27.0 SE). We retrieved 92 dances from the hives in the rural landscape with a mean foraging distance of 325.4 m (± 32.4 SE) and 73 dances from the hives in the urban landscape with a mean foraging distance of 465.8 m (± 43.8 SE). In the urban landscape foraging distances during lime tree flowering were significantly higher than foraging distances after lime tree flowering (mean foraging distance [m] ± SE: during 601.2 ± 57.6; after 260.4 ± 46.7). In the rural landscape foraging distances did not differ significantly during and after lime tree flowering (mean foraging distance [m] ± SE: during 269.3 ± 32.6; after 364.8 ± 49.9; LME results in Table II; pairwise comparisons in Supplement Table II; Fig. 2).

**Table II.** Linear mixed-effects model (LME) results of foraging distance in the rural and urban landscape during and after lime tree flowering. Pairwise comparisons in Supplement Table II.

|  | **Foraging distance [m]** |  |  |  |
| --- | --- | --- | --- | --- |
|  | Estimate | SE | t | p |
| Intercept | 304.81 | 59.32 | 5.14 | <0.001 |
| Urban | 119.19 | 86.79 | 1.37 | 0.303 |
| Rural:After | 69.85 | 70.26 | 0.99 | 0.344 |
| Urban:After | -338.81 | 103.97 | -3.26 | 0.009 |

**Fig. 2.**
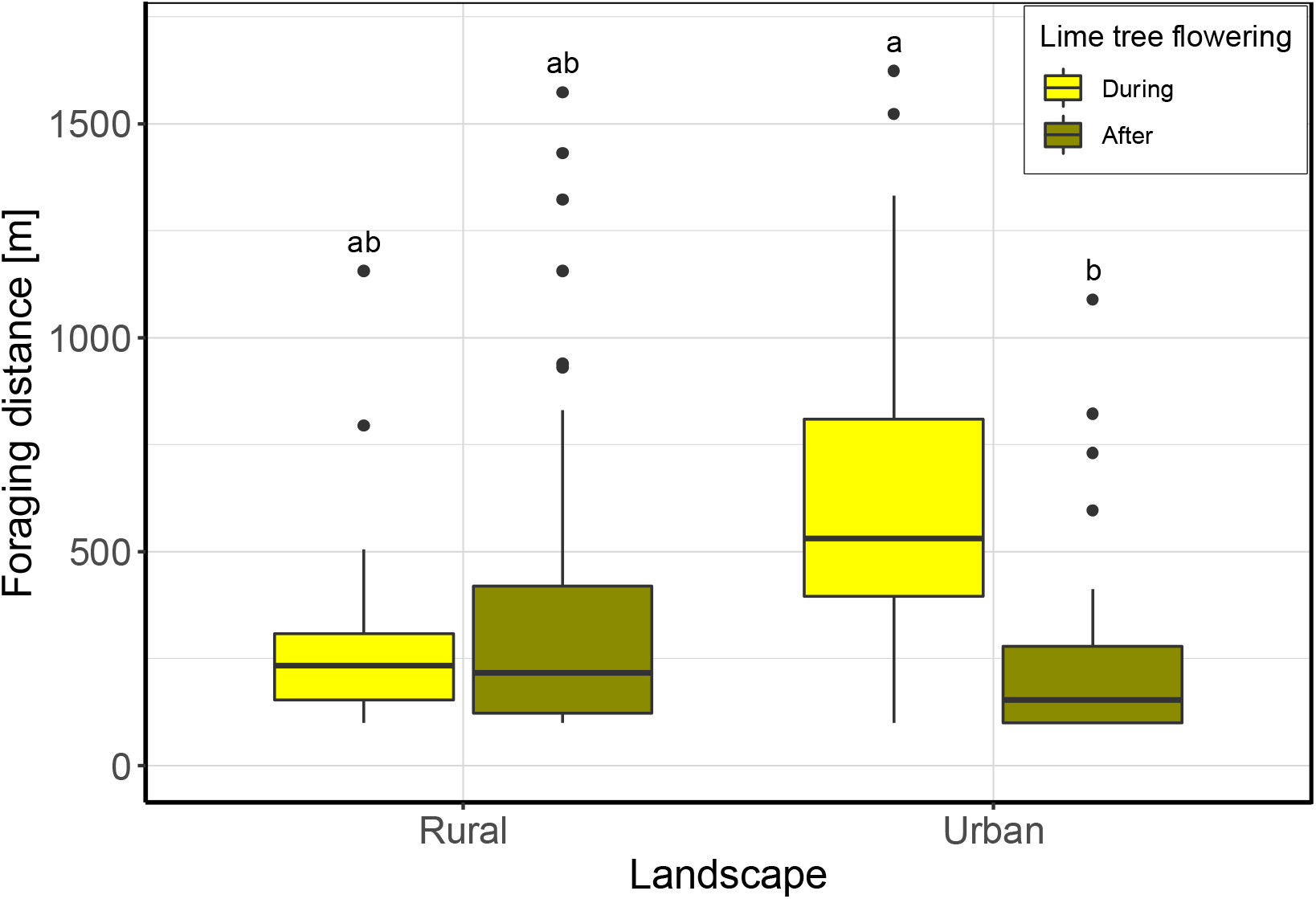
Foraging distance of honeybees in rural and urban landscapes during and after lime tree flowering. Letters over boxplots show differences between groups from Tukey’s pairwise post-hoc test (α = 0.05). LME results in Table II. Pairwise comparisons in Supplement Table II.

### Spatio-temporal variation in honeybee foraging habitats

Returning foragers mostly danced for foraging sites within a 1 km radius around the hives (urban: 7 of 73 dances, rural 5 of 92 dances outside of 1 km radius). In both the urban and the rural landscape foraging sites tended to be west of the hives during lime tree flowering. After lime tree flowering foraging sites were more evenly distributed in both landscapes. (Fig. 3).

**Fig. 3.**
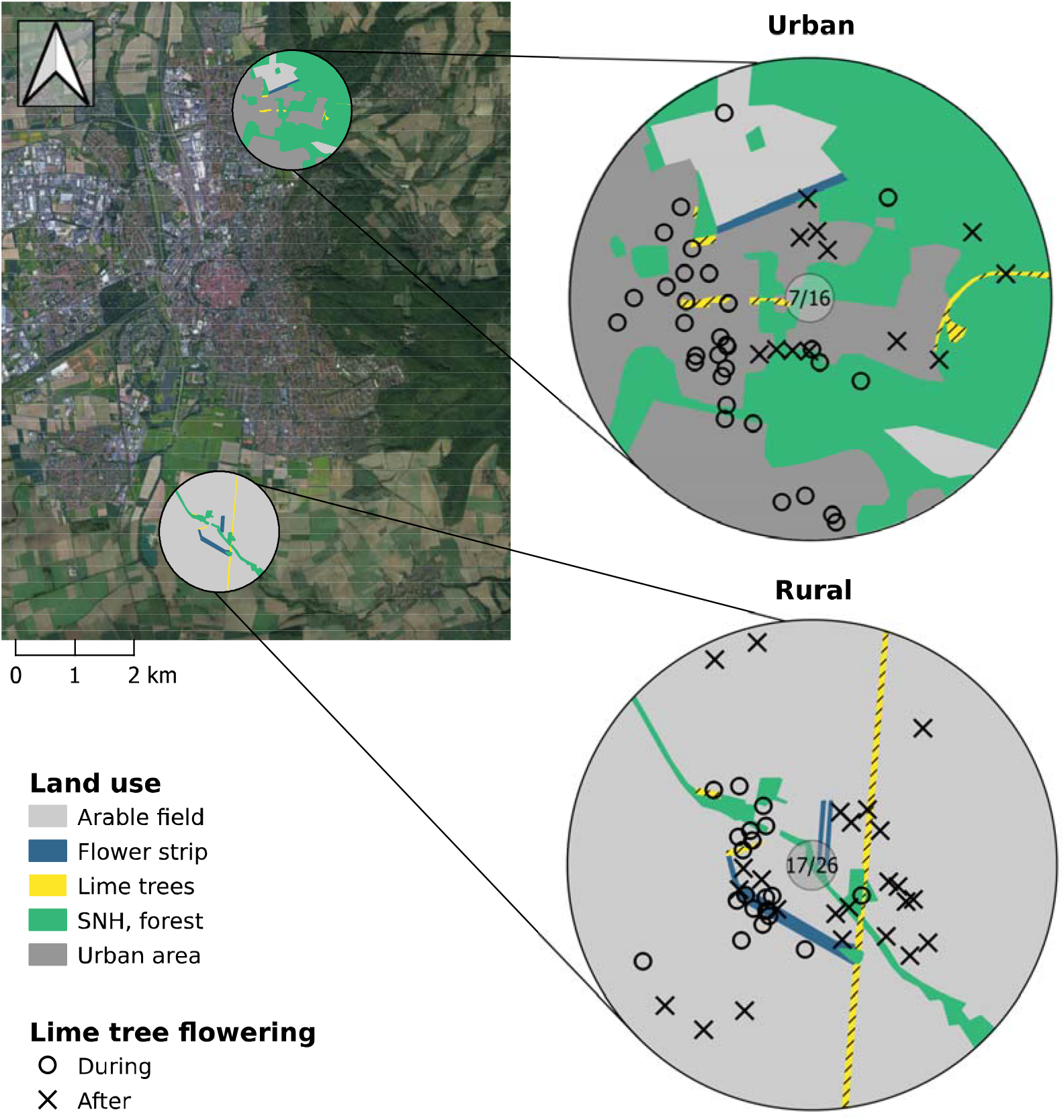
Spatial distribution of honeybee foraging habitats. Map shows the location of the hives in Göttingen, Germany. Spherical plots show land use in a 1 km radius around the honeybee hives in the urban and rural landscape, as well as foraging habitats during (open circles) and after (crosses) lime tree flowering. Numbers in the center of the spherical plots show the dances for local foraging sites before / after lime tree flowering.

### Pollen composition of pollen and nectar foragers

Overall, we counted 4136 pollen grains from 24 returning foragers during lime tree flowering (mean 199.6 pollen per pollen forager ± 0.4 SE; mean 145.1 pollen per nectar forager ± 22.5 SE). In the urban landscape 45.2% of the pollen of nectar-collecting honeybees came from lime trees and no pollen came from *P. tanacetifolia*. Pollen loads of pollen-collecting honeybees in the urban landscape were composed of 8.8% lime tree and 12.3% *P. tanacetifolia* pollen. In the rural landscape pollen of nectar-collecting honeybees was composed of 21.1% lime tree and 12.8% *P. tanacetifolia* pollen. Of the pollen of pollen-collecting honeybees in the rural landscape 10.8% came from lime trees and 53.0% came from *P. tanacetifolia* (Fig. 4).

**Fig. 4.**
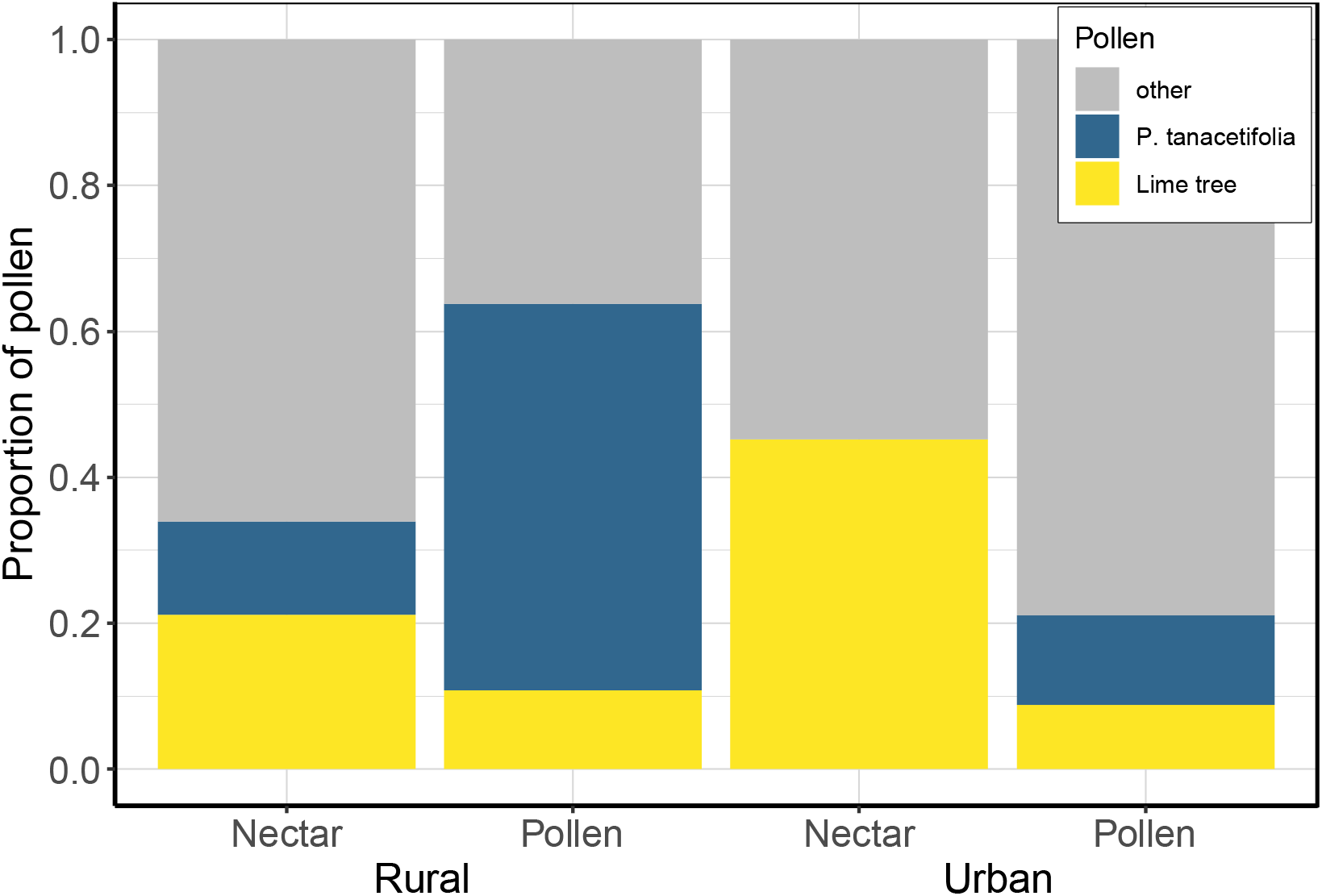
Proportion of *P. tanacetifolia* and lime tree pollen in honey stomachs of nectar-collecting honeybees and pollen from legs of pollen-collecting honeybees in the rural and urban landscape during lime tree flowering.

## 4. Discussion

Based on the combined analysis of bee abundance recordings, waggle dance decoding and analyses of pollen collected by nectar and pollen foragers, we demonstrate that foraging behaviour of honeybees depends on the spatio-temporal availability of highly rewarding resources. Our results furthermore indicate the relevance of lime trees as a nectar source in urban areas and *P. tanacetifolia* as a pollen source in rural areas.

Extended honeybee foraging ranges during lime tree flowering and large proportions of lime tree pollen in honey stomachs from nectar foragers in urban areas suggest that honeybees specifically adapt their foraging behaviour to exploit lime tree floral resources during this mass-flowering event. During lime tree flowering honeybees mainly foraged in the urban area west of the hives with aggregations of lime trees. Fewer foraging flights might have been directed at the lime trees east of the hives because those trees were smaller and might have provided less nectar than larger lime trees in the east of the hives (Kreider and Nehrkorn, pers. obs.). In rural areas, in contrast, foraging behaviour of honeybees was not affected by lime tree flowering to such an extent as in urban areas. Instead many foraging flights were directed at sown flower strips south-east of the hives which is mirrored in large proportions of pollen obtained from *P. tanacetifolia* by pollen foragers.

In contrast to previous studies (Steffan-Dewenter and Kuhn 2003; Danner et al. 2016; Danner et al. 2017; Bänsch et al. 2020) we found that honeybees did not extend their foraging range in response to floral resource scarcity but extended their foraging range to exploit a highly rewarding resource. This optimization of foraging behaviour in response to the availability of highly rewarding resources implies a trade-off between the distance travelled to a foraging site and the reward that is expected from the resource. This trade-off can be influenced by social organisation and communication of foraging sites if such decreases the energetic costs of foraging, i.e. the time spent searching for food (Cresswell et al. 2000; Lihoreau et al. 2011).

Plantation of lime trees in urban areas could effectively contribute to providing constant food supply for honeybees during summer. The extensive use of flower strips by honeybees and bumblebees in the rural landscape highlights that flower strips are an important flower substitute in otherwise flower poor agroecosystems. Especially, generalist bumblebees, such as *Bombus terrestris* L., can benefit from highly rewarding flower strips within their foraging range (Westphal et al. 2006a). However, our abundance counts of wild bees also indicate that wild bees barely benefit from flower strips as many wild bee species rely on particular plant species which typically do not occur in sown flower strips (Rollin et al. 2013). Consequently, different actions than only sowing flower strips need to be taken to conserve wild bees and protect the ecosystem services they provide than for the conservation of honeybees and bumblebees. These conservation actions include the preservation of SNH in rural areas and of flower-diverse garden habitats in urban areas which typically contain relatively high wild bee abundance and species diversity (Rollin et al. 2013; Martins et al. 2017), as well as the restoration of flower diversity in agroecosystems (Winfree 2010). Nevertheless, flowering trees, such as *Tilia* spp., *Acer* spp. L., *Robinia pseudoacacia* L. or *Aesculus hippocastanum* L. can play important roles for the provisioning of nectar and pollen to honeybees, bumblebees and some wild bee species in both rural and urban areas (Kämper et al. 2016).

During summer mass-flowering trees, such as lime trees, and sown flower strips are the most important foraging habitats of honeybees in urban and rural areas, respectively. Honeybees can adapt their foraging behaviour to spatial and temporal variation of floral resources in different foraging habitats and tend to optimize the exploitation of highly rewarding resources. In addition, our study shows that honeybees, bumblebees and wild bees exploit different foraging habitats within their foraging ranges which need to be restored and managed to conserve diverse wild bee communities and to provide sufficient floral resources to all managed honeybees, bumblebees and wild bees.

## Acknowledgements

We thank Susanne Jahn for managing the honeybee hives and Jan Schwedhelm for participating in the pollen identification. CW is grateful for funding by the Deutsche Forschungsgemeinschaft (DFG, German Research Foundation) – project number 405945293.

## Author contributions

CW designed the study. JJK and AN conducted the field recordings, decoded the waggle dances and determined bee abundances. CK conducted the pollen analysis. JJK and AN analysed the data with assistance from SB and CW. All authors contributed to the writing of the manuscript.

