## Supplementary Materials for "Honeybees optimize their foraging behaviour in relation to spatio-temporal changes in nectar and pollen availability"

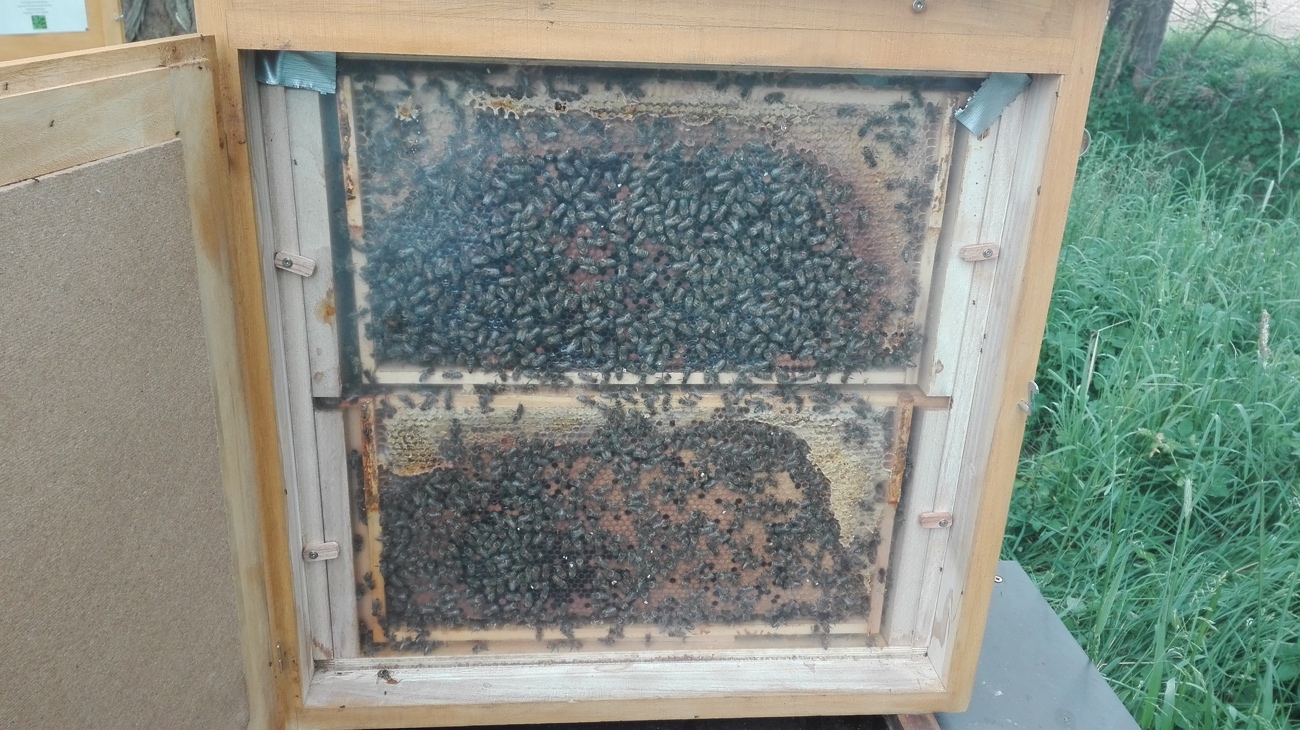

**Supplement Fig. 1** Honeybee observation hive as used in our study for waggle dance decoding.

**Supplement Table I** Tukey’s pairwise post-hoc test for generalized linear mixed model of square-root-transformed abundances of different functional bee groups (bumblebee, honeybee, wild bee) in different habitats (flower strip, SNH, urban area).

| Contrast | Estimate | SE | df | t-ratio | p |
| --- | --- | --- | --- | --- | --- |
| bumblebee, flower strip - honeybee, flower strip | 1.02 | 0.84 | 36 | 1.22 | 0.947 |
| bumblebee, flower strip - wild bee, flower strip | 8.72 | 0.84 | 36 | 10.38 | <0.001 |
| bumblebee, flower strip - bumblebee, SNH | 5.97 | 0.92 | 36 | 6.48 | <0.001 |
| bumblebee, flower strip - honeybee, SNH | 6.96 | 0.92 | 36 | 7.55 | <0.001 |
| bumblebee, flower strip - wild bee, SNH | 7.80 | 0.92 | 36 | 8.48 | <0.001 |
| bumblebee, flower strip - bumblebee, urban area | 6.35 | 0.99 | 36 | 6.44 | <0.001 |
| bumblebee, flower strip - honeybee, urban area | 5.89 | 0.99 | 36 | 5.98 | <0.001 |
| bumblebee, flower strip - wild bee, urban area | 8.48 | 0.99 | 36 | 8.60 | <0.001 |
| honeybee, flower strip - wild bee, flower strip | 7.70 | 0.84 | 36 | 9.16 | <0.001 |
| honeybee, flower strip - bumblebee, SNH | 4.95 | 0.92 | 36 | 5.37 | <0.001 |
| honeybee, flower strip - honeybee, SNH | 5.93 | 0.92 | 36 | 6.44 | <0.001 |
| honeybee, flower strip - wild bee, SNH | 6.78 | 0.92 | 36 | 7.36 | <0.001 |
| honeybee, flower strip - bumblebee, urban area | 5.33 | 0.99 | 36 | 5.40 | <0.001 |
| honeybee, flower strip - honeybee, urban area | 4.87 | 0.99 | 36 | 4.94 | <0.001 |
| honeybee, flower strip - wild bee, urban area | 7.45 | 0.99 | 36 | 7.56 | <0.001 |
| wild bee, flower strip - bumblebee, SNH | -2.75 | 0.92 | 36 | -2.99 | 0.101 |
| wild bee, flower strip - honeybee, SNH | -1.77 | 0.92 | 36 | -1.92 | 0.606 |
| wild bee, flower strip - wild bee, SNH | -0.92 | 0.92 | 36 | -1.00 | 0.984 |
| wild bee, flower strip - bumblebee, urban area | -2.37 | 0.99 | 36 | -2.41 | 0.311 |
| wild bee, flower strip - honeybee, urban area | -2.83 | 0.99 | 36 | -2.87 | 0.130 |
| wild bee, flower strip - wild bee, urban area | -0.25 | 0.99 | 36 | -0.25 | 1.000 |
| bumblebee, SNH - honeybee, SNH | 0.99 | 1.00 | 36 | 0.99 | 0.984 |
| bumblebee, SNH - wild bee, SNH | 1.83 | 1.00 | 36 | 1.84 | 0.655 |
| bumblebee, SNH - bumblebee, urban area | 0.38 | 1.06 | 36 | 0.36 | 1.000 |
| bumblebee, SNH - honeybee, urban area | -0.08 | 1.06 | 36 | -0.08 | 1.000 |
| bumblebee, SNH - wild bee, urban area | 2.51 | 1.06 | 36 | 2.38 | 0.327 |
| honeybee, SNH - wild bee, SNH | 0.85 | 1.00 | 36 | 0.85 | 0.994 |
| honeybee, SNH - bumblebee, urban area | -0.61 | 1.06 | 36 | -0.57 | 1.000 |
| honeybee, SNH - honeybee, urban area | -1.07 | 1.06 | 36 | -1.01 | 0.982 |
| honeybee, SNH - wild bee, urban area | 1.52 | 1.06 | 36 | 1.44 | 0.874 |
| wild bee, SNH - bumblebee, urban area | -1.45 | 1.06 | 36 | -1.38 | 0.899 |
| wild bee, SNH - honeybee, urban area | -1.91 | 1.06 | 36 | -1.81 | 0.673 |
| wild bee, SNH - wild. bee, urban area | 0.67 | 1.06 | 36 | 0.64 | 0.999 |
| bumblebee, urban area - honeybee, urban area | -0.46 | 1.11 | 36 | -0.41 | 1.000 |
| bumblebee, urban area - wild bee, urban area | 2.13 | 1.11 | 36 | 1.91 | 0.610 |
| honeybee, urban area - wild bee, urban area | 2.59 | 1.11 | 36 | 2.33 | 0.354 |

**Supplement Table II** Tukey’s pairwise post-hoc test for linear mixed-effects model of foraging distance in urban and rural landscapes during and after lime tree flowering.

| Contrast | Estimate | SE | df | t-ratio | p |
| --- | --- | --- | --- | --- | --- |
| Rural, After - Urban, After | 120.4 | 112.4 | 2 | 1.1 | 0.738 |
| Rural, After - Rural, During | 98.8 | 99.4 | 10 | 1.0 | 0.756 |
| Rural, After - Urban, During | -260.0 | 110.9 | 2 | -2.3 | 0.328 |
| Urban, After - Rural, During | -21.6 | 116.5 | 2 | -0.2 | 0.997 |
| Urban, After - Urban, During | -380.4 | 108.4 | 10 | -3.5 | 0.024 |
| Rural, During - Urban, During | -358.8 | 115.1 | 2 | -3.1 | 0.211 |
